# The human-specific *BOLA2* duplication modifies iron homeostasis and anemia predisposition in chromosome 16p11.2 autism patients

**DOI:** 10.1101/693952

**Authors:** Giuliana Giannuzzi, Paul J. Schmidt, Eleonora Porcu, Gilles Willemin, Katherine M. Munson, Xander Nuttle, Rachel Earl, Jacqueline Chrast, Kendra Hoekzema, Davide Risso, Katrin Männik, Pasquelena De Nittis, Ethan D. Baratz, 16p11.2 Consortium, Yann Herault, Xiang Gao, Caroline C. Philpott, Raphael A. Bernier, Zoltan Kutalik, Mark D. Fleming, Evan E. Eichler, Alexandre Reymond

## Abstract

Human-specific duplications at chromosome 16p11.2 mediate recurrent pathogenic 600 kbp BP4-BP5 copy number variations, one of the most common genetic causes of autism. These copy number polymorphic duplications are under positive selection and include 3–8 copies of *BOLA2*, a gene involved in the maturation of cytosolic iron-sulfur proteins. To investigate the potential advantage provided by the rapid expansion of *BOLA2*, we assessed hematological traits and anemia prevalence in 379,385 controls and individuals who have lost or gained copies of *BOLA2*: 89 chromosome 16p11.2 BP4-BP5 deletion and 56 reciprocal duplication carriers in the UK Biobank. We found that the 16p11.2 deletion is associated with anemia (18/89 carriers, 20%, *P*=4e-7, OR=5), particularly iron-deficiency anemia. We observed similar enrichments in two clinical 16p11.2 deletion cohorts, with 6/63 (10%) and 7/20 (35%) unrelated individuals with anemia, microcytosis, low serum iron, or low blood hemoglobin. Upon stratification by *BOLA2* copy number, we found an association between low *BOLA2* dosage and the above phenotypes (8/15 individuals with three copies, 53%, *P*=1e-4). In parallel, we analyzed hematological traits in mice carrying the 16p11.2 orthologous deletion or duplication, as well as *Bola2*^*+/-*^ and *Bola2*^*-/-*^ animals. The deletion and *Bola2*-deficient mice showed early evidence of iron deficiency, including a mild decrease in hemoglobin, lower plasma iron, microcytosis, and an increased red blood cell zinc protoporphyrin to heme ratio. Our results indicate that *BOLA2* participates in iron homeostasis *in vivo* and its expansion has a potential adaptive role in protecting against iron deficiency.

## Introduction

The human 16p11.2 chromosomal region is a hotspot of recurrent pathogenic copy number variants with diverse sizes, breakpoints, and gene content. Most breakpoints map within homologous segmental duplication clusters, consistent with non-allelic homologous recombination (NAHR) (Loviglio et al. 2017). Among these variants, 600 kbp deletions and duplications with breakpoints BP4 and BP5 are among the most frequent genetic causes of neurodevelopmental and psychiatric disorders (Weiss et al. 2008; McCarthy et al. 2009; Zufferey et al. 2012; D’Angelo et al. 2016; Marshall et al. 2017). They are also associated in a dose-dependent manner with head circumference, body mass index, age at menarche, and the size of brain structures associated with reward, language, and social cognition (Shinawi et al. 2010; Walters et al. 2010; Jacquemont et al. 2011; Maillard et al. 2015; D’Angelo et al. 2016; Martin-Brevet et al. 2018) (Männik et al., unpublished). BP4-BP5 rearrangements are mediated by *Homo sapiens sapiens*-specific duplications. These duplications appeared at the beginning of the modern human lineage, ∼282 thousand years ago, rapidly increased in frequency, and are now nearly fixed in humans, suggesting that a possible evolutionary advantage outweighs the accompanying chromosomal instability (Nuttle et al. 2016). The human-specific duplications at BP4 and BP5 contain three genes—*BOLA2, SLX1*, and *SULT1A*—that have single-copy orthologs in the mouse genome. Whereas *BOLA2* and *SLX1* have copies only in the BP4-BP5 flanking repeats, *SULT1A* has additional copies within the neighboring BP2. The BP4-BP5 duplicons also harbor copies of the primate gene family *NPIP* (Johnson et al. 2001), *BOLA2-SMG1* and *SLX1-SULT1A* fusion genes, and *SMG1P* and *LOC388242* pseudogenes (**Figure 1a**). These 102 kbp duplications are located at both ends of the single-copy region and are copy number variant in the genomes of contemporary humans (3 to 8 diploid copies) but are single copy in archaic genomes for which DNA sequence data are available (Nuttle et al. 2016).

**Figure 1.**
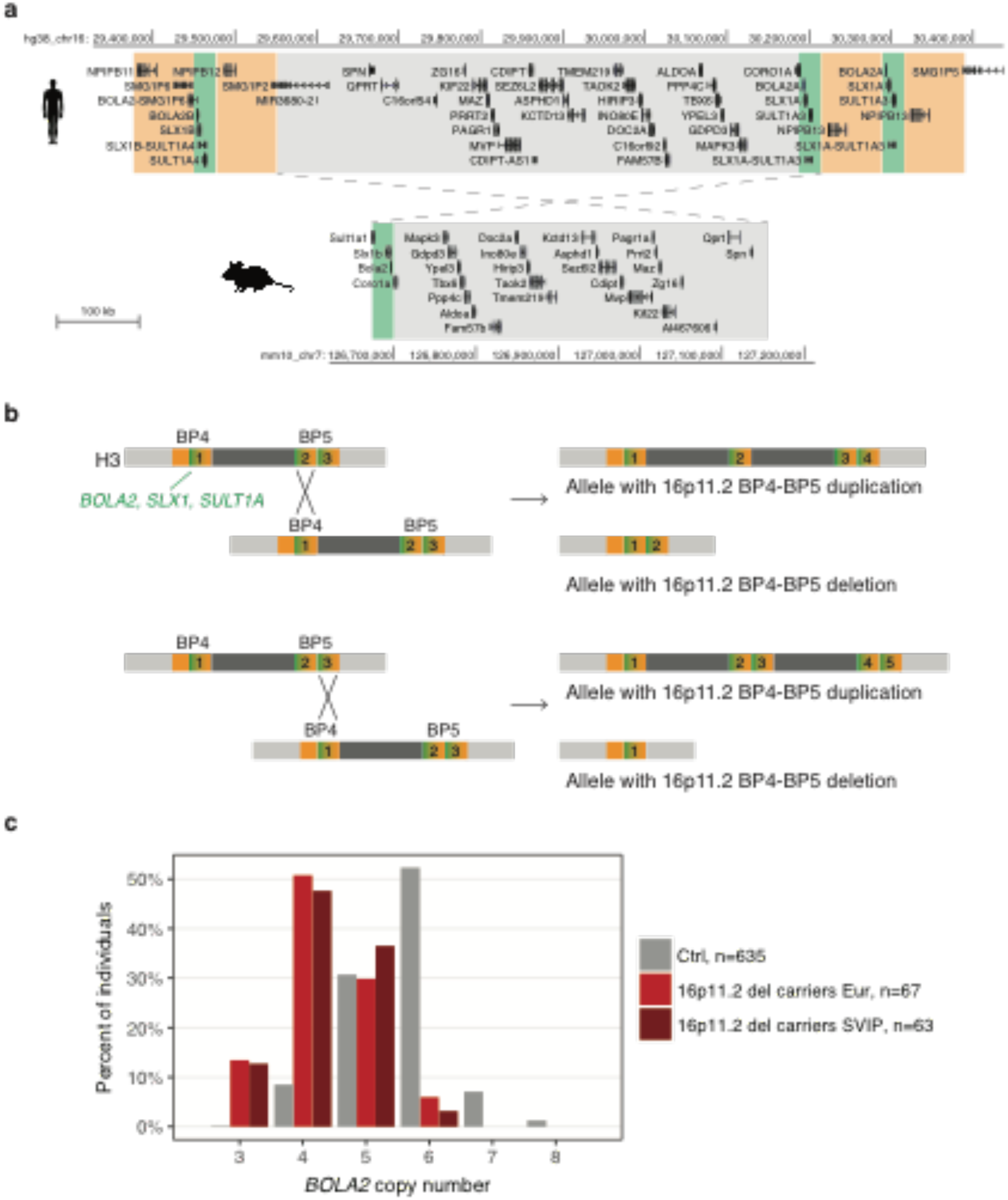
The 16p11.2 and *BOLA2* locus architecture. **a)** Gene content of the most common human haplotype of the 16p11.2 BP4-BP5 region (*top*) and comparison with the syntenic single-copy region rearranged in 16p11.2 CNV mouse models (Arbogast et al. 2016) (*bottom*). Human single-copy genes are grey-shaded; the *BOLA2*-*SLX1*-*SULT1A* human duplicon is green-shaded; the human-specific *NPIP* duplicon is orange-shaded. At BP4, the *BOLA2B, SLX1B*, and *SULT1A4* paralogs map; at BP5, two copies of the *BOLA2A, SLX1A*, and *SULT1A3* map. **b)** Two examples of outcomes from non-allelic homologous recombination between copies of the *BOLA2*-*SLX1*-*SULT1A-NPIP* segment at BP4 and BP5 are presented. The BP4-BP5 structure corresponds to the most common H3 haplotype. For example, sequence misalignments could occur between one BP4 copy and either the distal (*top* panel) or proximal (*bottom* panel) BP5 copies. The resulting alleles with a BP4-BP5 duplication and deletion have, respectively, more and fewer copies of *BOLA2, SLX1, SULT1A*, and *NPIP* (right panels). Numbers count the number of copies of the *BOLA2*-*SLX1*-*SULT1A* green segment in each haplotype. **C)** *BOLA2* copy number in 635 European individuals from the 1000 Genomes Project and Human Genome Diversity Project (Ctrl), 67 16p11.2 BP4-BP5 deletion carriers from the European cohort, and 63 deletion carriers from the SVIP cohort.

In this work, we investigated the potential adaptive role of the human-specific duplications at 16p11.2 BP4 and BP5. We focused our study on the possible benefits associated with an increased dosage of *BOLA2*, as this is the only gene in the segment with convincing evidence of duplications restricted to *Homo sapiens* (Nuttle et al. 2016). As *BOLA2* copy number positively correlates with both RNA expression and protein level in humans, and *BOLA2* expression in stem and lymphoblastoid cells is higher in human than in other large-bodied apes (Marchetto et al. 2013; Nuttle et al. 2016), we hypothesize that a possible advantage of *BOLA2* expansion derives from increased expression levels in some human cells. *BOLA2* encodes a protein that physically interacts with GLRX3 (glutaredoxin 3) to form a [2Fe-2S]-bridged heterotrimeric complex (Li et al. 2012). This complex participates in the maturation of cytosolic iron-sulfur proteins, transferring [2Fe-2S] clusters to CIAPIN1 (anamorsin), an essential component of the cytosolic iron-sulfur assembly (CIA) system (Zhang et al. 2008; Banci et al. 2015; Frey et al. 2016). In yeast, the [2Fe-2S]-bridged heterodimeric complex formed by the BolA-like protein Fra2 and the Grx3/Grx4 glutaredoxins has also essential roles in intracellular iron signaling and regulation (Kumanovics et al. 2008; Li and Outten 2012). GLRX3 and CIAPIN1 support erythropoiesis, as the knockdown of *glrx3* in zebrafish impairs heme synthesis and hemoglobin maturation (Haunhorst et al. 2013), and *Ciapin1-*deficient mouse embryos are anemic (Shibayama et al. 2004).

We investigated phenotypes associated with *BOLA2* copy number variation (CNV) in humans and mice and found that reduced *BOLA2* dosage associates with mild anemia in both species, suggesting a model wherein increases in *BOLA2* dosage might improve systemic iron homeostasis.

## Material and Methods

### Genotyping *BOLA2* copy number and statistical analysis in clinical cohorts

We estimated the *BOLA2A* and *BOLA2B* copy number of 16p11.2 BP4-BP5 deletion carriers with a molecular inversion probe (MIP) assay and probes mapping within *BOLA2* as described in (Nuttle et al. 2016). We collected iron metabolism-related (serum iron, ferritin, transferrin, coefficient of saturation of iron in transferrin), hematological parameters, and/or information about diagnosis of anemia for these individuals. For patients from the 16p11.2 Consortium, a diagnosis of anemia or microcytosis was established if the individual had low blood iron, low hemoglobin according to (World Health Organization 2011), MCV < 80 fl, or received a diagnosis of anemia. Within the SVIP cohort, a diagnosis of anemia was established if the individual was identified as having anemia per caregiver or self-report on the standardized medical history interview and confirmed through medical records review. We used Fisher’s exact test (R Development Core Team 2008) to assess the statistical significance of the difference in anemia prevalence between individuals with low and high number of *BOLA2* copies.

### CNV calling and association statistical analysis in UKB population cohort

UKB (Sudlow et al. 2015) is a volunteer-based general population biobank of the United Kingdom. Half a million participants aged 40-69 years at the time of recruitment (2006-2010) were enrolled through National Health Service patient registers. Participants consented to provide personal and health-related information, biological samples, and to have their DNA tested. The UKB governing Research Ethics Committee has approved a generic Research Tissue Bank approval to the UKB, which covers the research using this resource.

Single-nucleotide polymorphism genotyping in the UKB was performed using the UK BiLEVE and UKB Affymetrix Axiom platforms. Produced Log R ratio (LRR) and B allele frequency (BAF) values were formatted for CNV calling with hidden Markov model-based software PennCNV v1.0.4 (Wang et al. 2007). We retained samples that passed the post-processing quality-control parameters. To minimize the number of false positive findings, detected CNVs were stratified using a quality score (QS) (Mace et al. 2016). Only validated (QS>|0.8|) carriers were included in downstream analyses. We considered “16p11.2 BP4-BP5 CNV carriers” those that had a deletion or duplication starting in the interval 29.4-29.8 Mbp and ending 30.05-30.4 Mbp (hg19). We restricted the analysis to unrelated participants who declared themselves as white British.

We searched for evidence of association between gene dosage at 16p11.2 BP4-BP5 (deletion carriers, control individuals, and duplication carriers) and hematological traits relative to RBCs, reticulocytes (immature RBCs), and platelets, using linear models in the statistical package R (R Development Core Team 2008). We also ran pairwise comparisons of deletion carriers versus controls and duplication carriers versus controls using t-tests. In linear models, only additive effects for each copy (del=-1, controls=0, dup=1) were considered, and age, age^2, sex, the first 40 principal components from the genetic analysis, acquisition route, acquisition time, device ID, and freeze-thaw cycles were included as covariates. Trait measures were normalized by quantile transformation, prior to the analyses.

To assess the prevalence of anemia, we analyzed hospital discharge diagnoses coded following the International Statistical Classification of Diseases and Related Health Problems, 10th revision (ICD-10) together with self-declared illnesses. This information is available under data-fields 41202 and 41204 (respectively main and secondary diagnoses) and data-field 20002 of self-reported non-cancer medical conditions. We considered as anemic the individuals with ICD-10 codes D50, D51, D52, D53, D55, D58, D59, D60, D61, and D64, and self-reported non-cancer illness codes 1328, 1330, 1331, 1332, and 1446, as in (Crawford et al. 2019). A participant was coded as anemic if he/she had a diagnosis or self-reported the condition. We also used information under data-field 6179 (mineral and other dietary supplements) to assess the treatment with iron supplementation. We tested the difference in anemia prevalence between 16p11.2 deletion and duplication carriers versus controls using Fisher’s exact test.

### Mouse models

We used the 16p11.2^Del/+^ (Del/+) and 16p11.2^Dup/+^ (Dup/+) mouse models (Arbogast et al. 2016) and the C57BL/6N-Bola2^tm1(KOMP)Wtsi/Nju^ line (Bola2^tm1^) produced by the Knockout Mouse Project and International Mouse Phenotyping Consortium (Skarnes et al. 2011). Bola2^tm1^ animals were acquired from the Nanjing Biomedical Research Institute of Nanjing University, China.

For the Del/+, Dup/+, and Bola2^tm1^ Swiss cohort, all mice were born and housed in the Animal Facility of the Center for Integrative Genomics. We first removed the *agouti* allele from the Bola2^tm1^ line through backcrossing with C57BL/6N mice from Charles River. The animals were maintained on a standard chow diet (Kliba 3436, 250 ppm iron). All procedures were performed in accordance with protocols approved by the veterinary cantonal authority.

For the Bola2^tm1^ US cohort, heterozygous Bola2^tm1^ animals with the lacZ and neo cassettes were intercrossed to generate 8- and 15-week-old female animals for analysis. Furthermore, 129S4/SvJaeSor-*Gt(ROSA)26Sor*^*tm1(FLP1)Dym*^/J mice were acquired from the Jackson Laboratory and backcrossed (N>20) to the C57BL/6J background. The lacZ and neo cassettes in the Bola2^tm1^ animals were excised by breeding to the FLP1 mice and then backcrossed to C57BL/6J for an additional two generations. Resulting heterozygous animals were intercrossed to generate a cohort of 8-week-old female animals for analysis. All genetically modified mice were born and housed in the barrier facility at Boston Children’s Hospital. The animals were maintained on Prolab RMH 3000 diet (380 ppm iron; LabDiet). All procedures were performed in accordance with protocols approved by the Institutional Animal Care and Use Committee. In both animal facilities mice were provided water and food *ad libitum* and housed in 12-hour-long cycles of light and darkness.

### SMRT sequencing of *Bola2*^-/-^ mouse genome

Genomic DNA was isolated from the buffy coat layer of the blood of a male null mouse with the lacZ and neo cassettes excised (Kronenberg et al. 2018). We prepared one DNA fragment library (40-160 kbp inserts) with no additional shearing. After SMRTbell preparation, the library was size-selected with the BluePippin™ system (Sage Science) at a minimum fragment length cutoff of 40 kbp. Final library average size was 90 kbp. SMRT sequence data were generated using the PacBio Sequel instrument with Sequel Binding and Internal Ctrl Kit 3.0, Sequel Sequencing Kit 3.0, MagBead cleanup, diffusion loading, and acquisition times of 10-hour movies. A total of three SMRT Cell 1M v3 cells were processed yielding 11.5-fold (ROI/3.2 G) or 11.9-fold (raw/3.2G) whole-genome sequence data. The average subread length was 22.2 kbp with a median subread length of 14.1 kbp and N50 subread length of 35.9 kbp. Raw reads were mapped back to the mm10 mouse reference genome with minimap2 (Li 2018) and inspected manually at the *Bola2* locus and genome-wide with the PacBio structural variant calling and analysis tool. Local assembly of reads was done with the Canu assembler (Koren et al. 2017).

### Blood and tissue iron analysis

For the Del/+, Dup/+ and Bola2^tm1^ Swiss cohort, whole blood was collected from the tail vein in EDTA or heparin tubes (Sarstedt) for hematological and plasma iron measurements, respectively. Plasma was prepared by centrifugation for 10 minutes at 2,000 x g, 4 degrees. Measurements were executed by the Centre de PhénoGénomique, EPFL, Lausanne. The low body weight of 7-week-old Del/+ females did not allow collecting blood for hematological measurements due to ethical limits imposed by the veterinary office.

For the Bola2^tm1^ US cohorts, whole blood for complete blood counts was collected from the retro-orbital sinus into EDTA-coated microtainer tubes (Becton Dickinson) from animals anesthetized with ketamine/xylazine (100-120 mg/kg ketamine and 10 mg/kg xylazine) in sterile saline. Samples were analyzed on an Avida 120 analyzer (Bayer) in the Boston Children’s Hospital Department of Laboratory Medicine Clinical Core Laboratories. Whole blood for other purposes was collected by retro-orbital bleeding into serum separator tubes (Becton Dickinson), and serum was prepared according to the manufacturer’s instructions. Serum iron values were determined with the Serum Iron/TIBC kit (Pointe Scientific) according to the manufacturer’s instructions. Liver and spleen tissues were collected and tissue non-heme iron concentrations were determined as described previously (Torrance and Bothwell 1980). ZPP values in whole blood were analyzed on a ProtoFluor-Z Hematofluorometer (Helena Laboratories) according manufacturer’s instructions. Statistical analyses (t-test and linear model) were performed using the R software environment (R Development Core Team 2008).

## Results

### 16p11.2 deletion carriers have fewer copies of the *BOLA2*-*SLX1*-*SULT1A* segment

We previously sequenced haplotypes of the 16p11.2 BP4-BP5 interval (Nuttle et al. 2016). They differ in the number of copies and positions of a 102 kbp *BOLA2*-*SLX1*-*SULT1A* segment, i.e., its abundance within the BP4 or BP5 low-copy repeats. In particular, H3, the most common haplotype, has one copy of the segment at BP4 (including the paralogs *BOLA2B, SLX1B*, and *SULT1A4*) and two copies at BP5 (including the paralogs *BOLA2A, SLX1A*, and *SULT1A3*) (**Figure 1a**). As alleles with 16p11.2 BP4-BP5 CNVs are generated through NAHR between paralogous copies of this segment (Nuttle et al. 2016), we expect that deletion and duplication alleles would lose or gain copies of this segment along with one copy of the intervening unique region containing genes from *SPN* to *CORO1A* (**Figure 1b**). To assess this prediction, we quantified the number of copies of *BOLA2* in 16p11.2 BP4-BP5 deletion carriers collected by the 16p11.2 Consortium (n = 67, “European cohort”) and the Simons Variation in Individuals Project (SVIP, n = 63) (Nuttle et al. 2016).

Whereas European individuals from the 1000 Genomes Project and Human Genome Diversity Project (n = 635) have a mode of six copies of *BOLA2*, 16p11.2 deletion carriers showed a left-shifted distribution with a mode of four copies, confirming reduced *BOLA2* copy number (**Figure 1c**). Correspondingly, expression of *BOLA2* is positively associated with dosage in lymphoblastoid cells of 16p11.2 CNV carriers (Migliavacca et al. 2015). We identified nine and eight 16p11.2 deletion carriers with three *BOLA2* copies in the European (13.4%) and SVIP (12.7%) cohorts, respectively, compared to 0.2% (1 in 635) in control individuals. We never identified individuals with only two copies; one *BOLA2* copy per haploid genome is the ancestral copy number state observed both in great apes and archaic hominin genomes (Nuttle et al. 2016).

### Chromosome 16p11.2 microdeletion is associated with anemia and mild hematological defects

Since 16p11.2 BP4-BP5 CNVs affect the dosage of the copy number polymorphic *BOLA2, SLX1*, and *SULT1A* genes, it is possible that some associated phenotypes are due to the dosage change of one or a combination of these genes. Because *BOLA2* binding partners have been implicated in iron metabolism pathways and erythropoiesis, we evaluated hematological parameters, including red blood cell (RBC) indices and anemia, in 16p11.2 BP4-BP5 CNV carriers versus control individuals.

We examined CNVs in 488,366 individuals from the UK Biobank (UKB) and identified 89 unrelated Europeans (35 females and 54 males) carrying the 16p11.2 BP4-BP5 deletion and 56 (32 females and 24 males) carrying the duplication. We compared 18 hematological parameters to those of 379,385 control individuals with linear models and t-tests combining and separating the genders (**Supplemental Table S1**).

Gene dosage at 16p11.2 was negatively correlated with platelet count (PLTc, Beta = -0.685, *P* = 1e-15) and platelet mass per volume of blood (PLTcrit, Beta = -0.535, *P* = 4e-10), and positively associated with mean platelet volume (Beta = 0.444, *P* = 2e-7) (**Figure 2a**). Deletion carriers showed RBC anisocytosis (increased RBC distribution width, RDW, Beta = - 0.48, *P* = 2e-8) and a higher mean reticulocyte volume in males (*P* = 4e-4) (**Figure 2a**), but all other RBC traits were not different from controls.

**Figure 2.**
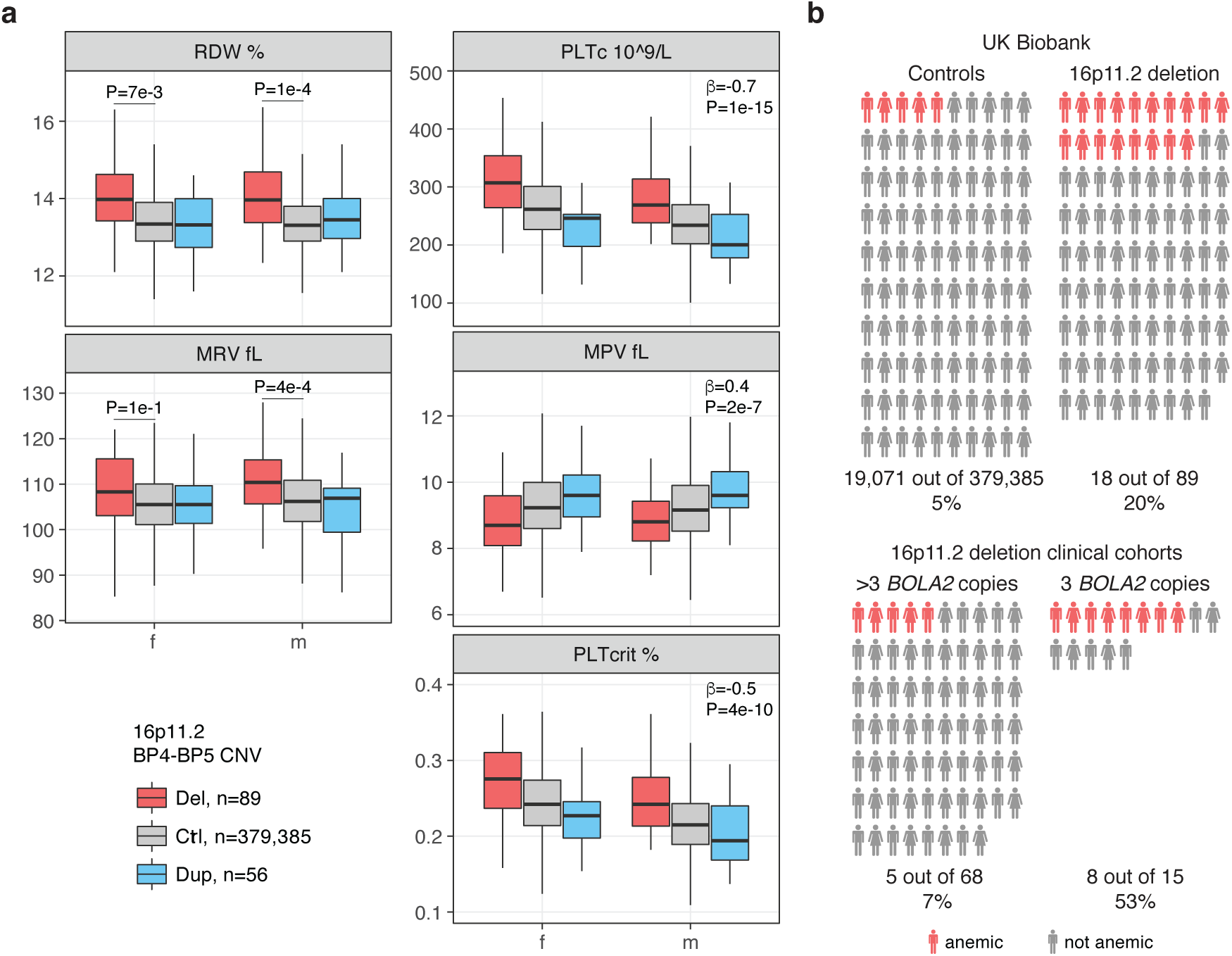
Hematological parameters and anemia prevalence in 16p11.2 BP4-BP5 CNV carriers. **a)** Red blood cell distribution width (RDW), mean reticulocyte volume (MRV), platelet count (PLTc), mean platelet volume (MPV), and platelet crit (PLTcrit) in 16p11.2 deletion and duplication carriers and control individuals of the UK Biobank (UKB). **b)** Prevalence of anemia in 379,385 control individuals and 89 16p11.2 deletion carriers of the UKB (*top*) and 83 16p11.2 deletion carriers of the SVIP and European cohorts stratified by *BOLA2* copy number (*bottom*).

We then tested a possible association between 16p11.2 gene dosage and the prevalence of anemia using both primary and secondary diagnoses collected for UKB participants. We found that the 16p11.2 deletion is associated with a higher prevalence of anemia (Fisher’s exact *P* = 4e-7, OR = 4.8) with 18 out of 89 carriers referred as anemic (20%) compared to 5% of control individuals (**Figure 2b**). Of these, four were females and 14 were males, with a trend towards higher prevalence of anemia in males (Fisher’s exact *P* = 0.1). Out of the 11 cases for which the type of anemia was specified, nine individuals had iron deficiency anemia (six with ICD-10 D50 diagnosis and three that self-reported the condition, three females and six males), one had vitamin B12 deficiency anemia, and one had autoimmune hemolytic anemia. We next assessed the prevalence of iron deficiency anemia (ICD-10 D50) in 16p11.2 deletion carriers versus control individuals and confirmed the higher prevalence in the former group (Fisher’s exact *P* = 4e-3). In contrast with deletion, 16p11.2 duplication carriers showed no association with anemia (Fisher’s exact *P* = 0.8).

### Reduced *BOLA2* copy number more strongly associates with anemia prevalence in 16p11.2 deletion carriers

Candidate genes for anemia in the 16p11.2 region are *BOLA2* and *ALDOA* (Frey et al. 2016). *ALDOA* missense mutations cause an autosomal recessive hemolytic anemia (Beutler et al. 1973), and this is distinctly different from the iron deficiency anemia associated with the 16p11.2 deletion. We thus interrogated the possible involvement of *BOLA2* and hypothesized that *BOLA2* CNV might contribute to the incomplete penetrance of anemia with the lowest copy number possibly conferring a higher risk. To test this hypothesis, we estimated *BOLA2* copy number and collected information regarding the presence of anemia, microcytosis, or iron deficiency in 83 families with the chromosome 16p11.2 deletion. This includes 63 deletion probands from the SVIP cohort (**Supplemental Table S2**) and 25 deletion carriers belonging to 20 families from the European cohort (**Supplemental Table S3**).

Six out of 63 probands from the SVIP cohort received a diagnosis of anemia (∼10%). We found a striking and significant difference in the prevalence of anemia between deletion carriers with three copies of *BOLA2* (4 out of 8, 50%) and those with more than three copies (2 out of 55, 4%) (Fisher’s exact *P* = 1.6e-3). In the European cohort, we collected partial hematological, ferritin, and iron measurements, or information on diagnosed medical conditions. Seven out of 20 (35%) unrelated deletion carriers had low hemoglobin concentration (mild or moderate anemia), low serum iron, and/or microcytosis, or received a diagnosis of anemia. Of these seven patients, four have three *BOLA2* copies and three have four. Here, we found a trend of higher anemia prevalence in individuals with three copies (Fisher’s exact *P* = 0.1) and observed that individuals both with three and four copies might be more susceptible to anemia, microcytosis, or iron deficiency (Fisher’s exact *P* = 0.08). We note that the European cohort subgroup for which we have phenotype data is enriched with individuals having three and four *BOLA2* copies. This might partly explain the different anemia prevalence in the two clinical cohorts (10% vs. 35%). Combining the two cohorts, the occurrence of anemia among deletion carriers with three *BOLA2* copies is significantly higher than that among those with more than three copies (Fisher’s exact *P* = 1e-4, OR = 13.6) (**Figure 2b**).

### Mouse models of the 16p11.2 deletion present with mild anemia and features indicative of early iron deficiency

To validate our human results, we assessed mouse models that carry a deletion (Del/+) or duplication (Dup/+) of the *Sult1a1*-*Spn* region syntenic to the human 16p11.2 BP4-BP5 locus (Arbogast et al. 2016). In these models, the rearranged region includes 26 genes that are in single copy in both the human and mouse genomes, together with the mouse single-copy *Sult1a1, Slx1b*, and *Bola2* whose orthologs map to the human-specific duplicated cassette. In contrast to other animal models of the 16p11.2 BP4-BP5 CNV (Horev et al. 2011; Portmann et al. 2014), these animals recapitulate the CNV both of the human single-copy genes and the copy-number variant *BOLA2, SLX1*, and *SULT1A* genes (**Figure 1a**). In particular, Del/+ mice allow studying the effects of reduced *Bola2* dosage in the context of the whole 16p11.2 BP4-BP5 rearrangement.

We measured plasma iron and 44 hematological parameters in Del/+ and Dup/+ male and female mice, as well as their respective wild-type littermates, at 7 (hematology only in males), 15, 29, and 50 weeks of age. In parallel, we recorded the body weight from 5 to 53 weeks of age (**Supplemental Figure S1**). As reported in (Arbogast et al. 2016), Del/+ male and female mice had a lower body weight. The weight of Dup/+ males was not different from that of their wild-type littermates, whereas Dup/+ females showed increased weight at 19 and between 23 to 29 weeks of age (*P* < 0.05). Del/+ mice showed lower plasma iron levels at 7, 15, 29, and 50 weeks of age and both genders. When we included both genotype and weight in the linear model, neither variable was significantly associated with iron level, probably because of the strong effect of the genotype on the weight in Del/+ mice. We observed no significant difference between Dup/+ mice and wild-type littermates although we noted a trend towards higher iron level in the former (**Figure 3a, Supplemental Table S4**). Some male mice had to be separated because of aggressive behavior. Their husbandry status affected the plasma iron, with male mice housed in single cages showing higher iron levels (*P* = 0.06 in the Del cohort at 50 weeks of age and *P* = 0.01 in the Dup cohort at 29 weeks of age) (**Supplemental Table S4**).

**Figure 3.**
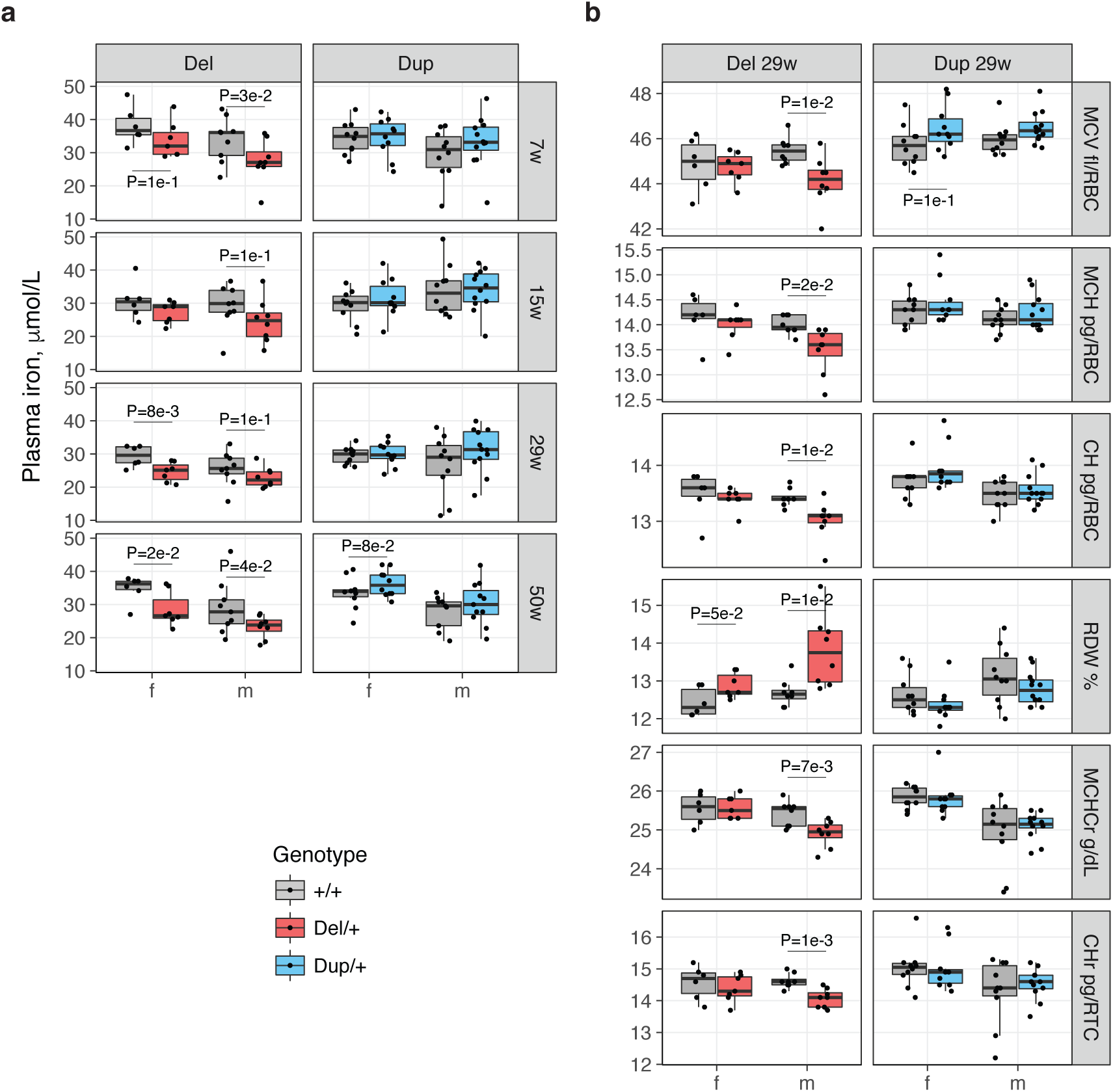
Blood traits of the 16p11.2^Del/+^ and 16p11.2^Dup/+^ mice. **a)** Plasma iron level in 16p11.2^Del/+^ and 16p11.2^Dup/+^ mice and their wild-type littermates at different ages (f: female, m: male, w: weeks). **b)** Red blood cell traits in 16p11.2^Del/+^ and 16p11.2^Dup/+^ mice and their wild-type littermates at the age of 29 weeks. These traits are significantly different in at least two longitudinal measurements (**Supplemental Tables S3 and S4**). *P* <= 0.1 are shown. (MCV: mean corpuscular volume, MCH: mean corpuscular hemoglobin, CH: red cell hemoglobin content, RDW: red cell distribution width, MCHCr: mean corpuscular hemoglobin concentration reticulocytes, CHr: red cell hemoglobin content reticulocytes.)

We describe here differences in hematological parameters between genotypes that differed significantly (*P* < 0.05) in at least two time points. Del/+ males showed lower mean corpuscular volume (MCV), mean corpuscular hemoglobin (MCH), red cell hemoglobin content (CH), mean corpuscular hemoglobin concentration of reticulocytes (MCHCr), and hemoglobin content of reticulocytes (CHr), and higher RDW, consistent with a mild hypochromic microcytosis (**Figure 3b, Supplemental Table S5**). Female mice showed a similar trend. No hematological parameter differed significantly between Dup/+ mice and wild-type littermates in more than one time replicate, in both females and males, except for a lower percentage of basophils in Dup/+ females (**Supplemental Table S6**).

### *Bola2* haploinsufficient and deficient mice have lower blood iron, smaller red blood cells, and higher zinc protoporhyrin and mean platelet mass

We assessed the role of *Bola2* in organismal iron metabolism and blood traits, analyzing mouse models with ablation of *Bola2*. To control for genetic background and environmental variability, we measured iron-related and hematological parameters in *Bola2*^-/-^ (homozygous mutants, ko/ko), *Bola2*^+/-^ (heterozygous mutants, ko/+), and *Bola2*^+/+^ mice (wild-type littermates, +/+). We created three different cohorts: i) a Bola2^tm1^ Swiss cohort; ii) a Bola2^tm1^ US “neo-in” cohort; and iii) a Bola2^tm1^ US cohort where we excised the LacZ and neomycin cassette to eliminate potential secondary effects of the antibiotic selection gene embedded within the locus.

We first assessed the engineered genotype by sequencing the genome of one neomycin cassette-excised *Bola2*^*-/-*^ male. We generated ∼12-fold sequence coverage using the long-read single-molecule, real-time (SMRT) sequencing platform and mapped and assembled long reads containing the *Bola2* locus. All sequence reads (9/9) confirmed the presence of a 563 bp deletion corresponding to exons 2-3 (**Supplemental Figure S2**), replaced by a 216 bp insertion, which includes one copy of the FRT recognition site. Western blotting of liver tissue from *Bola2*^-/-^, *Bola2*^+/-^, and wild-type mice (“neo-in” animals) confirmed the absence and reduced level of BOLA2 (**Supplemental Figure S3**).

To create the Bola2^tm1^ Swiss cohort, we set 25 ko/+ x ko/+ pairs that generated 21 +/+, 54 ko/+, and 26 ko/ko mice born within the same three-day interval from 17 crosses, suggesting no prenatal lethality (**Figure 4a**). Ko/+ and ko/ko mice showed no weight difference compared to wild-type littermates between 5 to 29 weeks of age, or in liver weight at 31 weeks (**Figure 4b and Supplemental Figure S4**). We tested for a possible association between *Bola2* dosage and plasma iron level at three different ages, 8, 17, and 22 weeks, in both genders. We observed significant positive association in females at 8 weeks (Beta = 3, *P* = 2e-3) and trend in males at 22 weeks (Beta = 4.3, *P* = 6e-2) (**Figure 4c, Supplemental Table S7**). When adjusting for weight and/or husbandry condition (males only, single versus shared cage), the association of iron level with gene dosage remained. Of note, animals isolated to a single cage show significantly higher plasma iron levels, as observed in the Del and Dup mouse cohorts. As a result, we constructed two linear models to estimate the effect of the number of *Bola2* copies on the iron level, assessing mice in single and shared cages separately. The effect of the genotype on the iron level was stronger and significant only in male mice housed in shared cages. However, the slopes of these two correlations did not differ significantly (*P* = 0.15).

**Figure 4.**
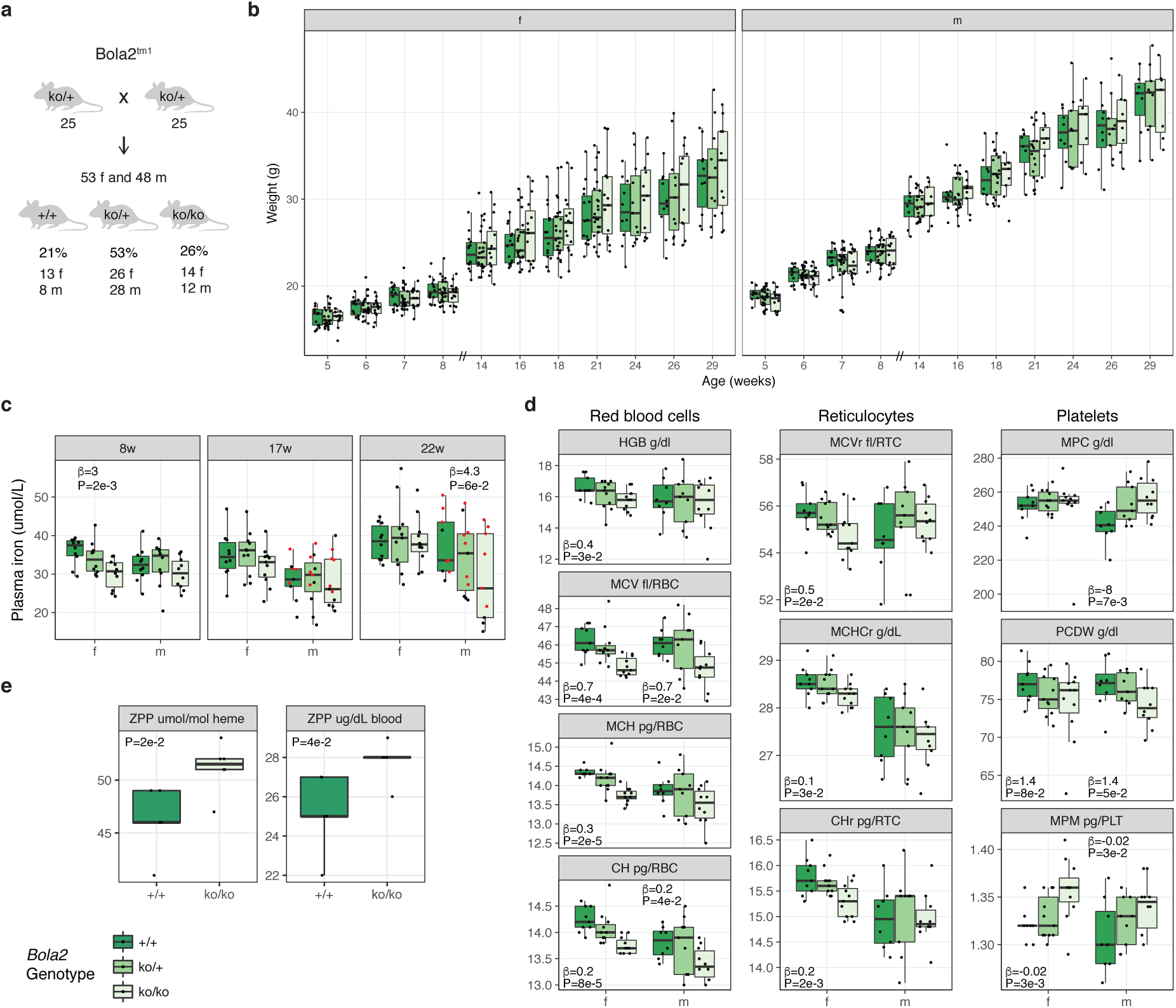
Characterization of the Bola2^tm1^ knockout mouse line. **a)** Mouse mating strategy of the Swiss Bola2^tm1^ line and gender and genotype ratios observed. **b)** Longitudinal body weight profiles. Note: the x-axis scale is not continuous. **c)** Plasma iron level in *Bola2*^+/-^ and *Bola2*^-/-^ mice and wild-type littermates (f: female, m: male, w: weeks). Red dots represent mice housed in single cages. **d)** Red blood cell (HGB: hemoglobin, MCV: mean corpuscular volume, MCH: mean corpuscular hemoglobin, CH: red cell hemoglobin content), reticulocyte (MCVr: mean corpuscular volume reticulocytes, MCHCr: mean corpuscular hemoglobin concentration reticulocytes, CHr: red cell hemoglobin content reticulocytes), and platelet (MPC: mean platelet component concentration, PCDW: platelet component distribution width, MPM: mean platelet mass) parameters of *Bola2*^+/-^ and *Bola2*^-/-^ mice and wild-type littermates at 17 weeks of age. Beta with *P* <= 0.1 are shown. **e)** Blood zinc protoporphyrin (ZPP) level of *Bola2*^-/-^ mice and wild-type littermates at 8 weeks of age.

*Bola2* dosage affects RBCs, reticulocytes, and platelets (**Figure 4d, Supplemental Table S8**). Specifically, the MCV (Beta = 0.7, *P*_males_ = 2e-2, *P*_females_ = 4e-4 at 17 weeks), MCH (Beta = 0.3, *P*_males_ = 2e-4 at 22 weeks, *P*_females_ = 2e-5 at 17 weeks), and CH (Beta = 0.2, *P*_males_ = 4e-2, *P*_females_ = 8e-5 at 17 weeks) decrease in *Bola2*^+/-^ and *Bola2*^-/-^ mice, in both genders. This shows that *Bola2* haploinsufficiency and deficiency are associated with a mild hypochromic microcytosis. Reticulocytes showed the same features, i.e., their size and hemoglobin content are positively associated with *Bola2* copy number. We observe that some platelet traits show differences: the PCDW (platelet component distribution width, Beta = 1.4, *P*_males_ = 5e-2, *P*_females_ = 8e-2) is positively associated with *Bola2* copy number, while the MPC (mean platelet component concentration, Beta_males_ = -8, *P*_males_ = 7e-3) and MPM (mean platelet mass, Beta = -0.02, *P*_males_ = 3e-2, *P*_females_ = 3e-3) are negatively associated.

To generate the US cohorts, heterozygous Bola2^tm1^ mice on a pure C57BL/6N background were intercrossed. Bola2^tm1^ animals with the LacZ and neo cassettes excised with the *Gt(ROSA)26Sor*^*tm1(FLP1)Dym*^/J FLP transgenic line on the C57BL/6J background were backcrossed to C57BL/6J for two more generations. Resulting heterozygous animals were intercrossed to yield (B6N)B6J N_3_F_1_ animals. We assessed complete blood counts and performed additional tests, including serum and tissue iron levels (**Supplemental Tables S9-12**). We replicated the mild microcytosis in the “neo-in” cohort (*P* = 1e-2) and observed no alteration in iron levels or spleen/body weight ratio suggestive of abnormal iron storage or an anemia compensated by erythroid hyperplasia. There were no morphological differences on peripheral blood smears. However, there were elevated zinc protoporhyrin (ZPP) normalized to heme levels in the blood of *Bola2*^-/-^ mice (*P* = 2e-2 in the “neo-in” and *P* = 1e-3 in the “neo-excised” cohort, **Figure 4e**), suggesting functional iron deficiency in maturing erythroid cells that could account for the mild microcytosis.

These data show that *Bola2* haploinsufficiency and deficiency cause mild hypochromic, microcytic, iron-deficiency anemia and suggest that a decrease in *BOLA2* dosage might be causative for the iron-deficiency anemia seen in 16p11.2 deletion carriers.

## Discussion

This work sheds light on some possible benefits of increased *BOLA2* dosage in humans, with six copies (the most common copy number), on average, per genome compared to two copies in other hominins, primates, and mammals (Nuttle et al. 2016). Duplications containing *BOLA2* map to the 16p11.2 locus and generate chromosomal instability associated with neurodevelopmental and psychiatric disorders. *BOLA2* duplications are thus another example of genetic trade-off as their potential gain comes with the cost of predisposition to pathogenic rearrangements (Weiss et al. 2008; McCarthy et al. 2009; Shinawi et al. 2010; Walters et al. 2010; Jacquemont et al. 2011; Zufferey et al. 2012; Maillard et al. 2015; D’Angelo et al. 2016; Nuttle et al. 2016; Marshall et al. 2017; Martin-Brevet et al. 2018), similarly to the positively selected 17q21.31 inversion associated with increased fecundity but predisposing to a deletion syndrome (Stefansson et al. 2005; Koolen et al. 2006).

We assessed blood-related phenotypes associated with *BOLA2* copy number and found that, as recently reported (Crawford et al. 2019), 16p11.2 BP4-BP5 deletion is associated with anemia (18/89, ∼20% of carriers), particularly iron deficiency anemia (9/11 cases with specified type of anemia), in a large adult population cohort, and men seemed to be more affected than women. In 16p11.2 deletion clinical cohorts, ∼50% of carriers with three *BOLA2* copies are anemic, suggesting a contribution of lower *BOLA2* dosage. We validated these results in mouse models of the 16p11.2 deletion, encompassing the *BOLA2* ortholog, and *Bola2*-specific haploinsufficiency and deficiency. All models presented iron deficiency and a mild form of microcytic and hypochromic anemia. Recalling the higher susceptibility of men, among Del/+ mice, only males had microcytosis, lower corpuscular hemoglobin, and anisocytosis. Transcriptome profiling of the brain cortex shows that Del/+ females but not males have increased expression of *Glrx3* and *Bola1*, thus possibly compensating the decreased *Bola2* expression through increased expression of BOLA2 binding partner (GLRX3) or mitochondrial counterpart (BOLA1). Finally, *Bola2* knockout mice showed mild microcytosis, higher ZPP level, and lower plasma iron, although the last phenotype was not observed in all longitudinal measurements possibly due to compensatory mechanisms. Our data suggest that *BOLA2* is a driver of iron deficiency anemia; however, it is still possible that other genes in the 16p11.2 single-copy region or its flanking human-specific duplicon contribute to this phenotype, as the number of *BOLA2* copies coincides with the number of copies of the entire 102 kbp segment. the effects of *BOLA2* copy number polymorphism on the phenotypic spectrum and incomplete penetrance of phenotypes of the 16p11.2 deletion is a striking example of the contribution of genes in the flanking low-copy repeats to traits associated with a genomic disorder, pointing out the evolutionary and medical relevance of this work.

We found discrepancies in hematological parameters between human and mouse. Both Del/+ male and *Bola2*^-/-^ and *Bola2*^+/-^ mice had significantly lower MCV and MCH compared to wild-type mice. We did not find the same features in RBCs of 16p11.2 deletion carriers possibly due to an unreported iron supplementation therapy for anemia. In fact, we found no difference in dietary iron supplementation between 16p11.2 deletion carriers and controls from the UKB (data-field 6179, comparing iron versus “none of the above”, Fisher’s exact *P* = 0.6). We note that this information might be partial as it was collected by a touchscreen questionnaire. Platelet traits were overtly modified in human and mouse; however, humans showed negative effects of gene dosage on PLTc and PLTcrit and positive effects on platelet volume, whereas Del/+ male, *Bola2*^-/-^ and *Bola2*^+/-^ mice showed higher platelet mass. These platelet alterations could be caused by lower dosage of *BOLA2* and/or other 16p gene/s or be secondary to the iron deficiency, as serum iron is inversely related to PLTc and PLTcrit in iron deficiency anemia (Dan 2005; Kadikoylu et al. 2006; Park et al. 2013). Human versus mouse discrepancies further confirm the fundamental divergence in gene expression between human and mouse hematopoiesis and the limitations of mouse systems to model human hematopoiesis (Pishesha et al. 2014; Beer and Eaves 2015).

Overall our data suggest that increased *BOLA2* dosage in humans might protect against iron deficiency. Iron deficiency is still the most common micronutrient deficiency in the world and the most common cause of anemia (Kassebaum and Collaborators 2016), suggesting the difficulty to precisely tune iron metabolism in humans. Iron is an essential micronutrient and serves as a cofactor for numerous enzymes, including those involved in energy metabolism, synthesis of DNA and proteins, and a range of other biochemical functions in cells. It is important in brain development, kidney function, immune responses, and growth and is essential for oxygen transport and delivery as component of hemoglobin and myoglobin (2012).

Veterinary hematological data for chimpanzee, bonobo, gorilla, and orangutan (Miller and Fowler 2015) (**Supplemental Table S13**) show that hemoglobin and iron levels in the blood do not differ between humans and great apes, suggesting that humans need higher BOLA2 levels to guarantee the same level of iron supply to the organism. Menstrual blood loss is much larger in human than in great apes. Different hypothesis have been proposed to explain this difference: elimination of pathogens from the reproductive tract; economy of energy as it is more costly to maintain the endometrium than to grow a new one; elimination of defective embryos; and adaptation to increased amounts of easily absorbable iron from meat (Denic and Agarwal 2007). It is possible that the increased copy number of *BOLA2* is necessary to cope with the increased menstrual iron loss, although males seemed to be more affected by anemia among 16p11.2 deletion carriers.

Higher BOLA2 levels might have been important in the past to thrive in nutrient poor environments or during the transition from a red meat hunter-gatherer Paleolithic diet to the iron-reduced cereal grain Neolithic diet, although *BOLA2* expansion is dated much earlier (Nuttle et al. 2016). These shifts likely resulted in an increased risk of iron deficiency anemia, especially in women of reproductive age. The rapid expansion and selection of *BOLA2* copy number at the root of the *Homo* lineage may have evolved to deal with this pressure as our species successfully expanded its ecological range at the cost of increased predisposition to rearrangements associated with autism.

## 16p11.2 Consortium members

Giuliana Giannuzzi, Katrin Männik, Catia Attanasio, Sandra Martin, Sébastien Jacquemont, Armand Bottani, Marion Gérard, Sacha Weber, Aurélia Jacquette, Fabien Lesne, Bertrand Isidor, Cédric Le Caignec, Mathilde Nizon, Catherine Vincent-Delorme, Brigitte Gilbert-Dussardier, Aurora Currò, Alessandra Renieri, Daniela Giachino, Alfredo Brusco, Alexandre Reymond. The affiliations and email addresses of the members of the 16p11.2 Consortium are listed in **Supplemental Table S14**.

## Acknowledgements

We thank families participating to the SVIP and the 16p11.2 Consortium cohorts. GG is recipient of a Pro-Women Scholarship from the Faculty of Biology and Medicine, University of Lausanne. This work was supported by grants from the Swiss National Science Foundation (31003A_160203 and 31003A_182632 to AR), National Institutes of Health (R01HG002385 to EEE), Horizon2020 Twinning project ePerMed (692145 to AR), Simons Foundation Autism Research Initiative (SFARI #303241 to EEE, #198677 to RAB, and #274424 to AR), INSERM, CNRS, University of Strasbourg (ANR-10-LABX-0030-INRT and ANR-10-INBS-07-03 PHENOMIN to YH), framework program Investissements d’Avenir (ANR-10-IDEX-0002-02 to YH). This research has been conducted using the UK Biobank resource (#16389). The funders had no role in study design, data collection and analysis, decision to publish, or preparation of the manuscript. EEE is an investigator of the Howard Hughes Medical Institute. We thank Jacques S. Beckmann for helpful discussions.

## Author Contributions

GG, PJS, MDF, EEE and AR designed the study and contributed to data interpretation. XN, KH and DR performed the MIP assays. GG, EP and ZK analyzed the UKB data. RE and RAB collected and analyzed phenotype information of SVIP probands. KM and 16p11.2 Consortium members provided phenotype information of European patients. GG analyzed phenotype data of European patients. YH and XG provided the mouse models. KMM performed SMRT sequencing and analyzed the data. EDB and CCP performed the western blot. GG, PJS, GW, JC and PDN performed experiments on mice. GG and PJS analyzed the mouse data. GG and AR wrote the manuscript with contributions from PJS, MDF and EEE. All authors read and approved the final manuscript.

