## Supplemental Information for "The human-specific *BOLA2* duplication modifies iron homeostasis and anemia predisposition in chromosome 16p11.2 autism patients"

### Supplemental Figures

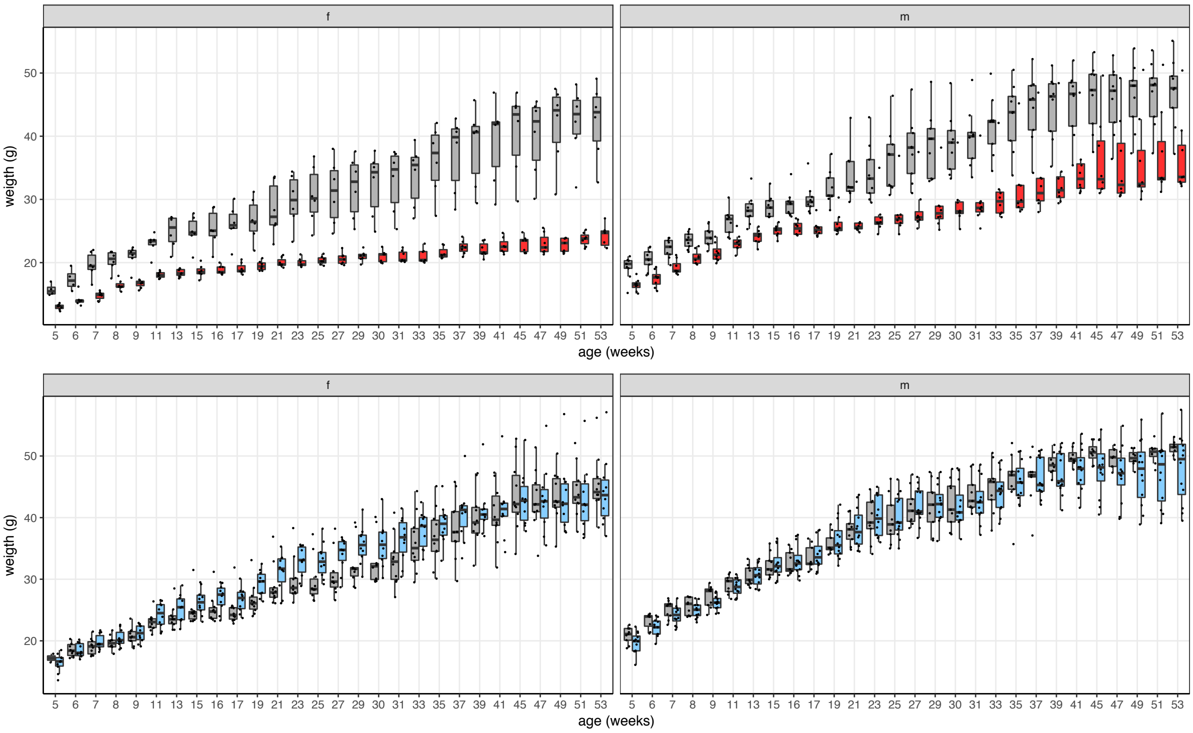


**Supplemental Figure S1.** Longitudinal body weight profiles of 16p11.2$^{Del/+}$ (*top*) and 16p11.2$^{Dup/+}$ (*bottom*) female (*left*) and male (*right*) mouse models and wild-type littermates. The genotype color code is: Del/+ (red), Dup/+ (light blue), wild-type (grey). Note that the x-axis scale is not continuous.

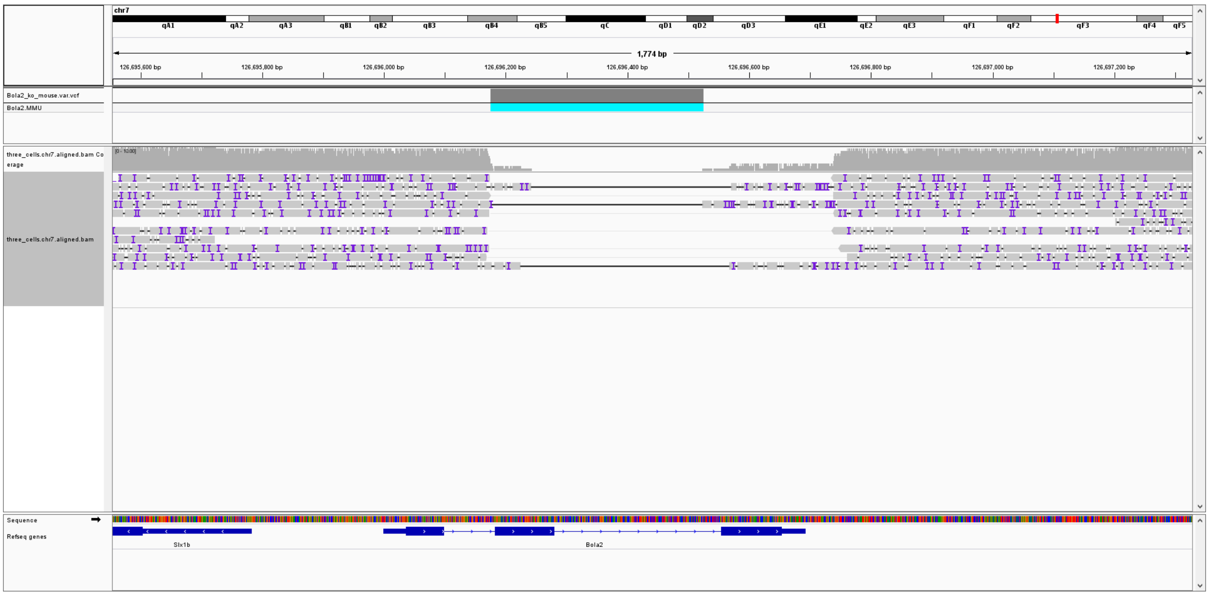


**Supplemental Figure S2.** SMRT sequencing platform reads of a neomycin cassette-excised *Bola2*$^{-/-}$ male were aligned to mm10. Within the *Bola2* locus on Chromosome 7 shown here please note the nine spanning reads, all indicating the deletion. PBSV calls a homozygous deletion of 351 bp (cyan box).

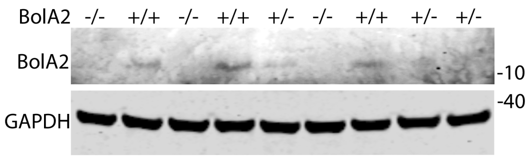


**Supplemental Figure S3.** Western blot of liver samples from *Bola2*$^{+/+}$, *Bola2*$^{+/-}$ and *Bola2*$^{-/-}$ mice with an anti-BOLA2 antibody performed as described in [17].

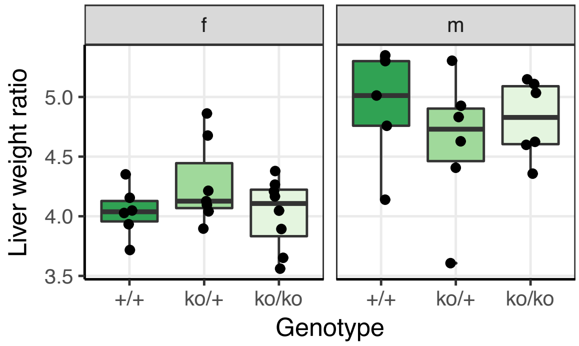


**Supplemental Figure S4.** Liver weight normalized to body weight (mg/g) of *Bola2*$^{-/-}$, *Bola2*$^{+/-}$ and wild-type littermates from the Swiss cohort at 31 weeks of age.

#### Supplemental Tables

**Supplemental Table S1.** Hematological parameters of 16p11.2 CNV carriers and controls of the UK Biobank.

**Supplemental Table S2**. *BOLA2* copy number and anemia status of 16p11.2 deletion carriers from the SVIP cohort.

**Supplemental Table S3.** Hematological and blood iron quantifications of 16p11.2 deletion carriers from the European cohort with genotyped *BOLA2* copy number. Values in red are below the minimum value of the reference range. Individuals not considered in the statistics are grey-shaded (related family members).

**Supplemental Table S4.** Heparin-plasma iron level of 16p11.2$^{Del/+}$ and 16p11.2$^{Dup/+}$ mouse models and their wild-type littermates.

**Supplemental Table S5**. Hematological parameters of 16p11.2$^{Del/+}$ mouse models and their wild-type littermates.

**Supplemental Table S6**. Hematological parameters of 16p11.2$^{Dup/+}$ mouse models and their wild-type littermates.

**Supplemental Table S7**. Heparin-plasma iron level of *Bola2*$^{+/+}$, *Bola2*$^{+/-}$, and *Bola2*$^{-/-}$ mice (European cohort).

**Supplemental Table S8**. Hematological parameters of *Bola2*$^{+/+}$, *Bola2*$^{+/-}$, and *Bola2*$^{-/-}$ mice (European cohort).

**Supplemental Table S9.** Serum iron, TIBC, Tf sat, ZPP, liver and spleen iron levels of *Bola2*$^{+/+}$, *Bola2*$^{+/-}$, and *Bola2*$^{-/-}$ mice (neo-in mice, US cohort).

**Supplemental Table S10.** Hematological parameters of *Bola2*$^{+/+}$, *Bola2*$^{+/-}$, and *Bola2*$^{-/-}$ mice (neo-in mice, US cohort).

**Supplemental Table S11.** Serum iron, TIBC, Tf sat, and ZPP levels of *Bola2*$^{+/+}$ and *Bola2*$^{-/-}$ mice (neo-excised mice, US cohort).

**Supplemental Table S12.** Hematological parameters of *Bola2*$^{+/+}$ and *Bola2*$^{-/-}$ mice (neo-excised mice, US cohort).

**Supplemental Table S13.** Normal hemoglobin and iron levels in the blood of adult humans, great apes, and mice.

**Supplemental Table S14.** Members of the 16p11.2 Consortium.
